# Parallel, Continuous Monitoring and Quantification of Programmed Cell Death in Plant Tissue

**DOI:** 10.1101/2023.08.22.554256

**Authors:** Alexander Silva Pinto Collins, Hasan Kurt, Cian Duggan, Yasin Cotur, Philip Coatsworth, Atharv Naik, Matti Kaisti, Tolga Bozkurt, Firat Güder

**Author notes:** These authors contributed equally to this work (Hasan Kurt, Cian Duggan, Yasin Cotur).

## Abstract

The accurate quantification of hypersensitive response (HR) programmed cell death is imperative for understanding plant defense mechanisms and developing disease-resistant crop varieties. In this study, we report an accelerated phenotyping platform for the continuous-time, rapid and quantitative assessment of HR: Parallel Automated Spectroscopy Tool for Electrolyte Leakage (PASTEL). Compared to traditional HR assays, PASTEL significantly improves temporal resolution and has high sensitivity, facilitating the detection of microscopic levels of cell death. We validated PASTEL by transiently expressing the effector protein AVRblb2 in transgenic lines of the model plant *Nicotiana benthamiana* (expressing the corresponding resistance protein Rpi-blb2) to reliably induce HR. We were able to detect cell death at microscopic intensities, where leaf tissue appeared healthy to the naked eye one week after infiltration. PASTEL produces large amounts of frequency domain impedance data captured continuously (sub-seconds to minutes). Using this data, we developed a supervised machine learning models for classification of HR. We were able to classify input data (inclusive of our entire tested concentration range) as HR-positive or negative with 84.1% mean accuracy (F_1_ score = 0.75) at 1 hour and with 87.8% mean accuracy (F_1_ score = 0.81) at 22 hours. With PASTEL and the ML models produced in this work, it is possible to phenotype disease resistance in plants in hours instead of days to weeks.

## Introduction

Plants depend on innate immunity to eliminate infectious agents and parasites. ^1^ This relies on effective recognition of pathogens leading to activation of intracellular signaling cascades to elicit defense responses. Cell-surface receptors have the capability to identify pathogen-associated molecular patterns (PAMPs) and pathogen-secreted proteins, commonly referred to as effectors, located in the extracellular space. In contrast, effector proteins translocated inside the host cells are typically recognized by intracellular nucleotide-binding domain leucine-rich repeat-containing proteins (NLRs)^2^. Activation of cell-surface immune receptors or NLRs often culminates in the hypersensitive response (HR), typified by a rapid and localized form of programmed cell death at the site of infection.^3,4^ HR can be generated in response to various classes of pathogens and is associated with disease resistance. On the macroscopic scale, HR cell death often manifests as discolored, necrotic lesions with abrupt boundaries from surrounding healthy tissue. HR is associated with numerous processes, including cytoplasmic shrinkage, loss of turgor pressure, and vacuolization^5,6^. The phenotype and kinetics of HR can, however, differ greatly depending on the host and causative agent.^7^ HR assays are invaluable in the study of plant-pathogen interactions,^8,9^ defense responses, and associated signal transduction pathways.^10^ HR assays are used not only for detecting the presence of HR, but also for precisely evaluating the intensity and timing of responses. Furthermore, HR assays can be used to screen for candidate gene/effector pairs,^11,12^ which confer disease resistance, critical to the development and engineering of more resistant crop varieties.

Assessment of HR is non-standardized and numerous methods are used to evaluate it, both qualitatively and quantitatively. In its simplest form, HR is assessed manually by visually inspecting the symptoms on the leaf surface with the naked eye and by applying an arbitrary index scoring system. This system rates the intensity based on the size and appearance of the lesions. Azo dyes such as Trypan blue and Evans blue penetrate non-viable cells and are commonly used to visualize cell death;^13,14^ microscopy imaging is performed on stained tissue, and cell death intensity can be estimated using image processing techniques. Staining can also be used to detect hydrogen peroxide, a reactive oxygen species produced during HR, using diaminobenzidine.^15^ Fluorescence imaging can be used to visualize the accumulation of HR-related phenolic compounds^16^ or detect changes in autofluorescence of leaves^17^ as a measure of cell death. Another technique for measuring HR is the electrolyte leakage assay (ELA), which involves placing excised leaf tissue in a small volume of distilled water and regularly sampling the conductivity of the solution.^18–20^ HR cell death compromises the integrity of cellular membranes, resulting in a decrease in the impedance of the sample solution as ionic species diffuse from the plant tissue into the sample. The extent of leakage is dependent on the intensity of cell death and so enables a quantitative proxy measure of HR.

While different methods have their own advantages and are often used in conjunction with each other as confirmatory measures of HR, few are suitable for high throughput, sensitive and continuous measurement.^21,22^ Staining techniques are typically both time and labor-intensive, can involve the handling of toxic chemicals, and each stained tissue sample provides data for only a single time point. High throughput image acquisition has been demonstrated with fluorescence imaging of whole leaves,^17^ but as with other optical techniques, quantification/scoring is not fully automated and requires manual processing. ELAs require minimal sample preparation and enable a quantitative readout at multiple time points but involve a large time commitment; conductivity is sampled manually, and so can be arduous if measuring many samples at regular (typically hourly) intervals.

A highly sensitive, automated system would offer several benefits for detecting the activation of HR in plant immunity. One significant advantage would be the ability to identify microscopic HR, a function that most conventional methods lack. This capability is crucial because certain conditions, although presumed not to cause HR, might indeed have microscopic HR occurring. Without such sensitive detection, these conditions could potentially lead to misleading conclusions. Moreover, an automated system can aid in studying the early events taking place before visual HR symptoms become apparent. This allows for a deeper understanding of the initial stages of HR activation, which can be pivotal in understanding plant immunity and developing prevention strategies against pests and diseases.

In this work, we describe a high-throughput ELA platform for rapid quantification of HR cell death. The proof-of-concept platform (**P**arallel **A**utomated **S**pectroscopy **T**ool for **E**lectrolyte **L**eakage, PASTEL) is implemented in an eight-channel modular well format with integrated and removable low-cost single-use electrodes alongside low-cost microcontroller-based electronics. PASTEL offers continuous time monitoring of electrolyte leakage through parallel automated impedance spectroscopy measurements with a tunable frequency range and adjustable time intervals (sub-seconds to minutes). Compared to traditional ELAs, the continuous time, high frequency sampling capabilities of PASTEL enables faster detection of the onset of HR and higher temporal resolution for changes in conductivity. We demonstrate the ability of PASTEL to quantify differing intensities of cell death, including detection of microscopic HR, using agroinfiltration mediated HR in transgenic lines of *Nicotiana benthamiana* with integrated controls.

## Results

### Multi-sample, automated conductivity measurement system

PASTEL consists of two main components: i) a modular, 3D-printed array of eight reusable sample measurement wells with replaceable, single-use conductivity probes and ii) a microcontroller-based system for sampling solution impedance. The measurement wells (**Figure 1a**) can each hold a set of three leaf discs (the sample) and analyte solution, with the electrode assembly inserted in the bottom aperture. The design with the electrodes placed at the bottom ensures that the floating tissue samples do not come into contact with the electrodes, interfering with the measurement. The electrodes are, therefore, only in contact with the analyte solution under test. The internal diameter of the wells (20 mm) is sufficient for each leaf disc (diameter of each leaf disc = 7.8 mm) to be completely in contact with the solution without overlapping with enough space for placing the discs in the desired (abaxial side down) orientation.^23^

**Figure 1.**
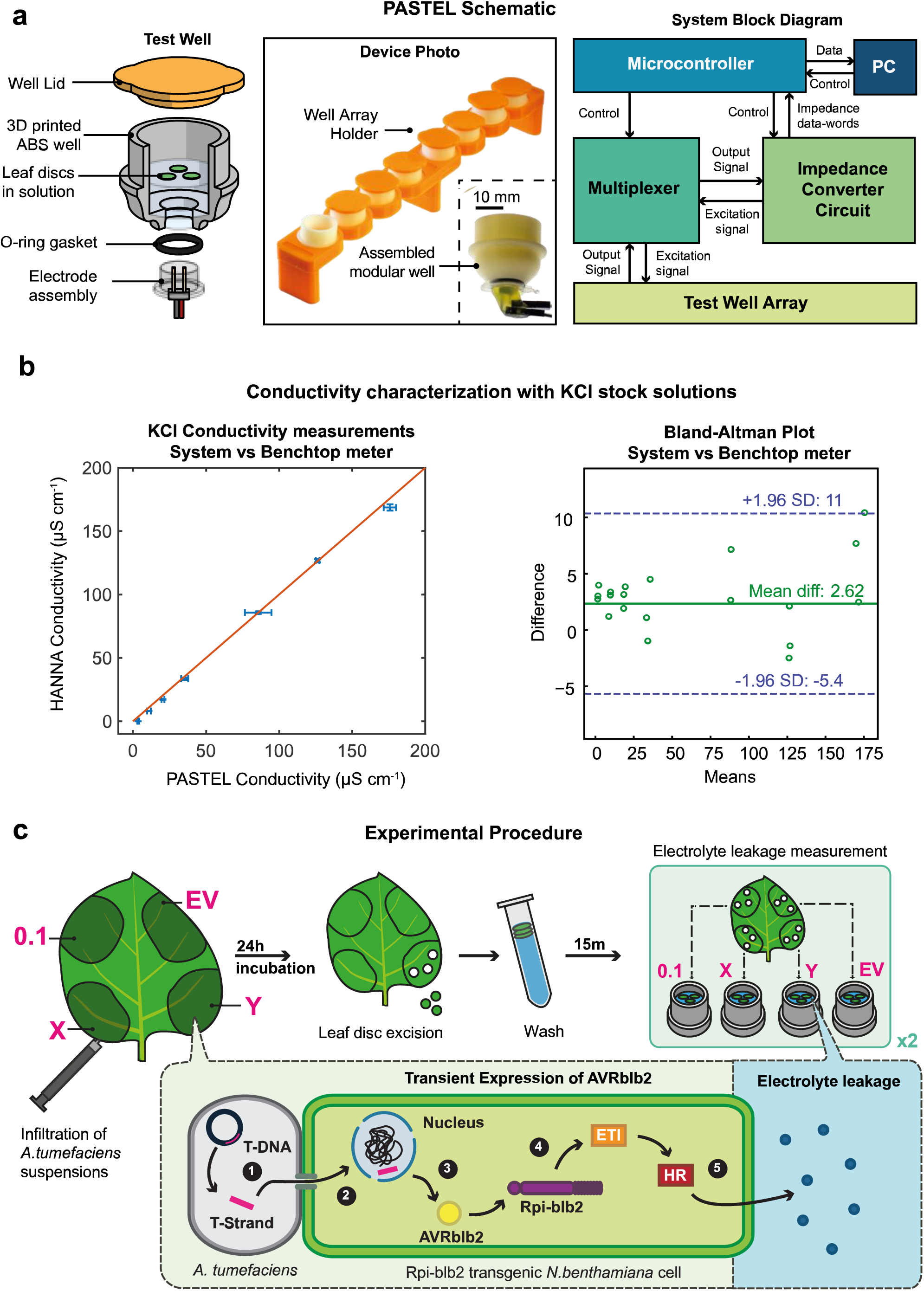
**a**) Exploded view schematic of individual measurement well assembly during measurement, the photograph of well assembly and system block diagram outlining key components and control hierarchy. **b**)Correlation graph of the commercial HANNA conductivity meter and PASTEL system while measuring KCl solutions ranging from 0 to 1 mM measured by (10 kHz excitation frequency, 198 mV_p-p_) (Data represented as µ ± σ, with n = 3 independent samples). Bland-Altman graph of PASTEL system against commercial HANNA conductivity meter in the specified KCl solutions. (±1.96 σ upper and lower limits designate the confidence level of 95%. **c**) Schematic of experimental procedure for leaf disc assay. OD_600_ = 0.1 and EV infiltration patches are the positive and negative controls, respectively. X and Y are discrete concentrations of agrobacterium suspension varied between experiments. Inset: Simplified molecular schematic of HR generation via agroinfiltration. 1. Single-stranded T-DNA (ssT-DNA) containing gene for expression for AVRblb2 is processed from the tumor inducing plasmid in the agrobacterium. 2. ssT-DNA is transported to the plant cell nucleus via a type IV secretion system. 3. Gene within ssT-DNA is expressed and AVRblb2 effector protein is synthesized. 4. AVRblb2 is specifically recognized by Rpi-blb2 NLR present in the transgenic plant, ETI response is initiated. 5. ETI culminates in HR, resulting in loss of cell membrane integrity and leakage of electrolytes into solution surrounding the leaf disc.

We designed the internal geometry of the well to minimize the experimental volume of liquid required while still providing sufficient surface area for multiple leaf discs to float without overlapping. A smaller sampling volume results in less dilution of the sample (reducing sensitivity requirements of the conductivity sensor) and reduced ionic diffusion time toward electrodes. We 3D-printed the wells using a filament of acrylonitrile butadiene styrene (ABS), a durable, water-resistant polymer, making them suitable for reuse following sterilization with ethanol. The sensor assembly is a simple two-electrode setup constructed from chemically inert gold-plated brass pins mounted in a polypropylene housing. The jumper wires were secured onto the housing by adhesive and connected the respective electrodes to the electronics. Nitrile rubber O-rings were used to create a water-tight seal between the electrode assembly and measurement well. To complete the entire assembly, the lids for the wells (used to minimize analyte evaporation and block ambient light) and a stand for the measurement wells were 3D printed using polylactic acid (PLA).

The electronics that perform the measurements facilitate continuous, concurrent acquisition of the solution impedance in each of the eight measurement wells. An AD5933 impedance converter (Analog Devices) integrated on a custom printed circuit board (PCB) was implemented to measure complex impedance over a frequency range of 5 – 100 kHz. By sampling the response signal, the AD5933 generates real and imaginary data to determine the magnitude and phase of the measured impedance, following software calibration. The input/output voltage pins of AD5933 were connected to the electrode assemblies via an 8-channel multiplexer (74HC4051, SparkFun) such that the excitation signal can be applied and programmatically switched between wells to perform near-simultaneous, parallel measurements. The measurement time for all 8 wells was less than 1 s at a single frequency and approximately 20 s for the entire 5 – 100 kHz frequency range (1 kHz intervals). An ATmega328P CH340 Nano microcontroller was used for the overall control of the measurement system (**Figure 1a**), interfacing with both the multiplexer and AD5933 via I^2^C. The measurement system was connected to a nearby PC over USB, for exchange of experimental data and commands between. The USB connection also provided all the electrical power needed to operate the measurement system.

Using an aqueous solution of KCl, ranging from 0 to 2 mM, we also calibrated the measurement system using an excitation voltage of 198 mV_P-P_ at 10 kHz. The solution conductivity was between 0 – 350 μS cm^-1^ in this range of concentrations. For validation, we also performed measurements using a commercially available reference device: HI-5521 Bench Meter (HANNA Instruments). By applying a scaling constant calculated using the reference measurements made with the 84 μS cm^-1^ conductivity standard (500 μM KCl), we were able to convert the magnitude of the conductance measured by PASTEL at 10 kHz into a conductivity value (**Equation S1**). Both the commercial device and PASTEL also produced highly linear measurements across the range of concentrations tested. Applying the scaling factor across the range of measurements taken at 10 kHz produced a strong agreement with the commercial instrument by HANNA (**Figure 1b**) with R^2^ = 0.9986. The Bland-Altman plot shows the mean difference in measured conductivity between PASTEL and the commercial device is small (2.62) but variance appears to increase with the conductivity of solution being measured. This systematic shift indicates that the system requires further optimization for higher ion concentrations or a more complex calibration function. In any case, the difference between measurements were within two standard deviations of each other. Calibration and characterization to determine impedance and phase measured over the entire frequency range was also performed (see **Figures S1, S2** and Methods).

### Quantitative detection of hypersensitive response with *Nicotiana benthamiana*

We used agroinfiltration as a means to induce a reliable, repeatable HR to test electrolyte leakage with our system (**Figure 1c**). Agroinfiltration is a technique that uses *Agrobacterium tumefaciens* as a vector to deliver a gene of interest into plant cells, where it is then transiently expressed. This gene of interest is housed within the transfer DNA (T-DNA), replacing a set of genes originally found in the tumor-inducing (Ti) plasmid. ^19^ Subsequently, the single-stranded T-DNA is processed and transferred to the plant cell nucleus via a type-IV secretion system, enabling the gene to be expressed without being incorporated into the genome.^24^

We used transgenic lines of the model plant *N. benthamiana* expressing the resistance (R) protein Rpi-blb2 in all experiments where HR was induced. Rpi-blb2 is a nucleotide-binding leucine-rich repeat (NLR) protein from *Solanum bulbocastanum* that senses the AVRblb2 family of effectors from the potato blight pathogen *Phytophthora infestans.*^24–26^ Upon specific recognition of AVRblb2, Rpi-blb2 initiates an effector-triggered-immunity (ETI) response that culminates in HR programmed cell death^27^ In each experiment, Rpi-blb2 *N. benthamiana* leaves were agroinfiltrated with suspensions of *A. tumefaciens* carrying AVRblb2 constructs in order to transiently express AVRblb2 and subsequently generate HR. Suspensions of *A. tumefaciens* carrying empty vector (EV) constructs were used for negative control infiltrations. The infiltration suspensions were quantified by optical density at the wavelength of 600 nm (OD_600_) upon initial preparation and subsequently diluted to the required OD_600_ (See Methods).

We conducted a series of experiments using a range of infiltration levels with a view to evaluate the key aspects of our system: i) reliable, binary detection of HR cell death vs. a negative control; ii) the ability to resolve different intensities of cell death and iii) the minimum run-time required for statistically significant detection of HR.

In each experiment, two leaves of a single Rpi-blb2 *N. benthamiana* plant were identically inoculated with a set of four *Agrobacterium* suspensions (**Figure 1c**). In each quadrant of the leaf, a different suspension was infiltrated; an empty vector-carrying negative control (OD_600_ = 0.1), two inocula X and Y, each containing AVRblb2-carrying bacterial suspensions in the test range of concentrations OD_600_ = 0.0005 – 0.05, and an AVRBlb2-carrying positive control with a relatively higher concentration (OD_600_ = 0.1) resulting in robust visual HR symptoms. The inocula X and Y were prepared at the desired concentrations by combining AVRblb2-carrying and EV-carrying suspensions such that an overall OD_600_ of 0.1 was achieved in order to control for any plant response caused by the bacteria itself. The plant was then incubated for 24 hours in a growth chamber before taking sets of three leaf discs from each infiltration patch (two replicates per infiltration, eight sets total per experiment). A representative leaf at the time of excision is shown in **Figure 2a**. Each set of discs was washed and placed in an individual measurement well with distilled water, abaxial side down. Three leaf discs were used per well in an attempt to account for any heterogeneity in expression, cell death within an infiltration patch, and to increase the analytical signal. Impedance measurements were recorded by the system over a 22-hour period, sampling over 5 – 100 kHz (with 1 kHz intervals) every two minutes.

**Figure 2.**
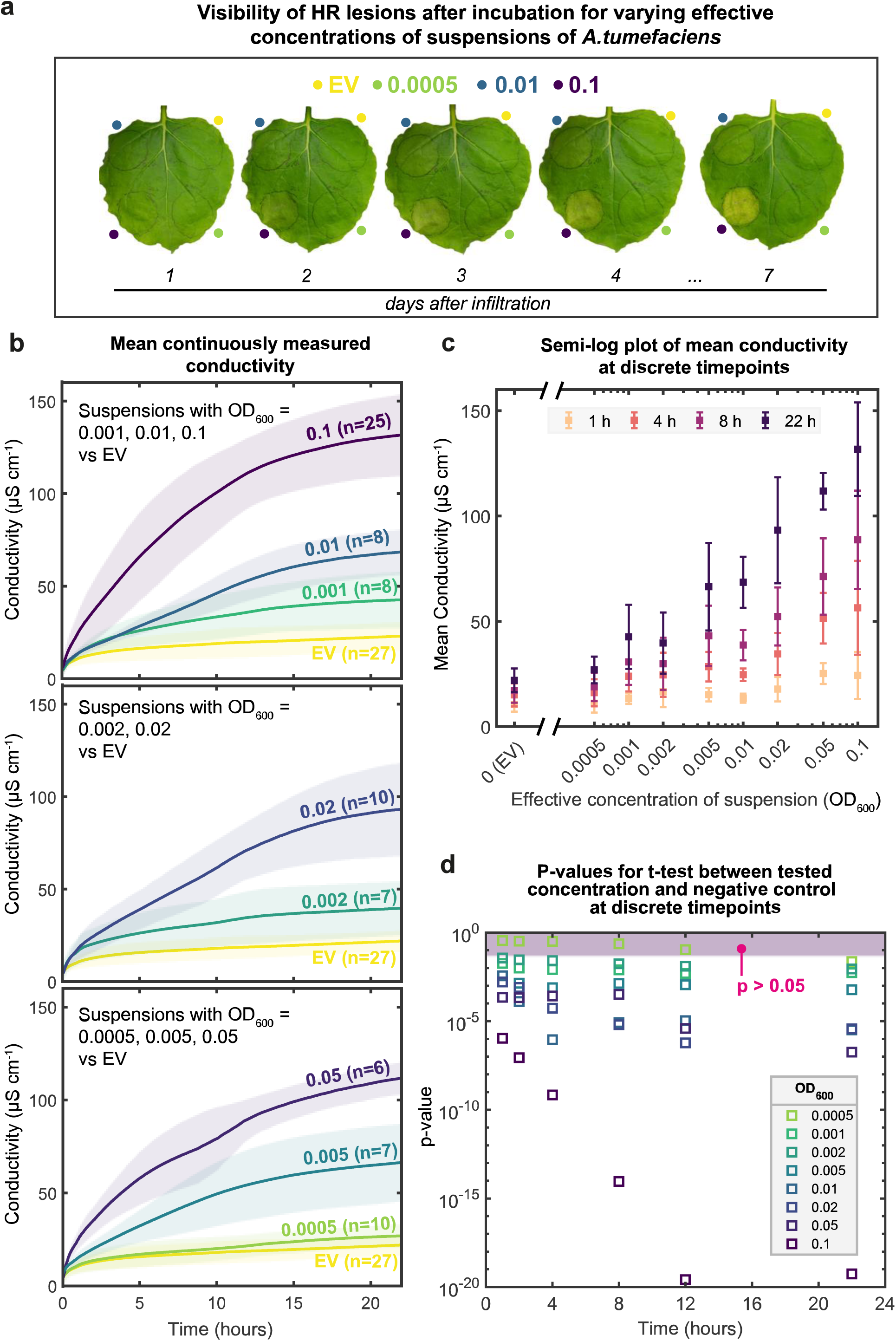
**a**) Images of a leaf agroinfiltrated with an empty vector-carrying bacterial suspension (negative control) and AVRblb2-carrying bacterial suspensions at concentrations of OD_600_ = 0.0005, 0.01 and 0.1. At 1 day (when leaf discs are excised when conducting experiments with PASTEL), all regions of infiltration appeared healthy. At 2 days, cell death was visible only in the region where the high positive control was infiltrated (OD_600_ = 0.1). The region where the low concentration (OD_600_ = 0.0005) was infiltrated remained indistinguishable from the negative control at 7 days. **b)** Mean solution conductivities measured over a 22-hour period for samples infiltrated with a range of concentrations of agrobacterium suspensions, averaged over multiple independent experiments. Trace labels denote the OD_600_ value of *Agrobacterium* suspension and number of repeats. Plots grouped by concentrations differing by an order of magnitude with control for reference. Data was represented as µ ± σ (shaded region), captured at 10 kHz excitation frequency, 2-minute sampling interval, moving average filter applied. **c)** Mean solution conductivities versus OD_600_ concentration of infiltration suspension at 1, 4, 8 and 22 hour time points. Same source data as in b). Data was represented as µ ± σ. **d)** p-values for a right-tailed, independent t-test testing the hypothesis that the mean solution conductivity for samples infiltrated with concentration z (denoted in legend) is greater than the mean solution conductivity of negative controls at time point x. The shaded region represents 5% significance level.

We were able to reliably detect HR with PASTEL for all nine infiltrations tested; mean solution conductivity (averaged at each time point across all repeat experiments, 10 kHz excitation frequency) for all infiltration concentrations tested was greater than that of the negative control within 1 hour of measurement. By the 22-hour endpoint, the mean conductivity of all but the lowest concentration exceeded the negative control by >80% (**Figure 2b**). The mean solution conductivity of the infiltration of OD_600_ = 0.0005 was only 22% greater than the negative control at the endpoint. The mean conductivity measured followed a similar progression for all concentrations tested; an initial rapid increase when the discs are first added to the solution, gradually decreasing towards a plateau at the endpoint where the concentration of the electrolytes within the plant tissue approaches equilibrium with the solution. The equilibrium conductivity is, of course, dependent on the intensity of cell death. This conductivity progression was seen even in the absence of hypersensitive response in the negative control, owing to a basal level of electrolyte leakage, likely caused by the trauma from the excision.

Comparing infiltrations differing in concentration by an order of magnitude or greater, clear differences in measured conductivity were seen within and often before 5 hours. Although concentrations less than an order of magnitude apart are distinguishable, the high variance would suggest that individually comparing data of similar concentrations across independent experiments may not be viable. Instead, multiple repeats should be performed to obtain reliable averages for comparison. We speculate a number of possible contributing factors to this high variance, including differences in plant age, variability in expression and inherent error in spectrophotometer measurements when preparing dilutions.

We would expect, in general, for expression of AVRblb2 to increase with increasing concentration of *Agrobacterium,* and so to the intensity of cell death. We observed a strong positive correlation between mean solution conductivity and concentration of infiltration, shown at time points of 1, 4, 8, and 22 hours (**Figure 2c**). As electrolytes permeate from the plant tissue, the measured conductivity differences increase, corresponding to varying intensities of cell death. Should complete cellular membrane deterioration occur, it would result in maximal endpoint solution conductivity. The logarithmic scaling suggests that this limit might be approached at the highest concentration tested. In preliminary experiments, agroinfiltration of AVRblb2-carrying suspensions of OD_600_ > 0.1 was attempted, but the degradation of plant tissue at these high concentrations was too severe, making it challenging to reliably extract intact leaf discs.

We found a statistically significant difference in mean solution conductivity compared to the negative control (right-tailed independent t-tests, *p* = 0.05) for all but the infiltration of OD_600_ = 0.0005 at the 1-hour time point (**Figure 2d**). With PASTEL, we were, therefore, able to rapidly detect HR for even low intensities of cell death. Statistical significance from the control was not found for the infiltration of OD_600_ = 0.0005 until the endpoint, but it should be noted that at this concentration, cell death is only present at the microscopic level; plant tissue appears healthy to the naked eye even several days following infiltration (**Figure 2a**).

### High-resolution sampling using the PASTEL system detects significant variance in plant cell death intensity and kinetics across different leaves

The rapid sampling rate of the system and ability to measure over a range of frequencies generate a large dataset for each channel that enables us to analyze individual experiments in more depth than with a traditional electrolyte leakage assay, where the sampling intervals are typically on the order of hours.^18^ Despite being treated identically, we observed marked differences in solution conductivity between two leaves of the same plant for the same infiltration concentrations, with the exception of the negative controls (**Figure 3a**). For each leaf in isolation, we observe the expected trend of endpoint conductivity increasing with higher infiltration concentration, but comparing across the leaves, we notice much higher endpoint conductivity for the infiltrations of OD_600_ = 0.1 and 0.02 in leaf B than in leaf A. Notably, the infiltration of OD_600_ = 0.02 in leaf B results in a higher endpoint conductivity than the infiltration of OD_600_ = 0.1 in leaf A. As previously reported^28^, this finding suggests that differences in agroexpression and resulting cell death intensity can be significant across different leaves of the same plant, likely a large source of the variance we observed when finding the mean conductivity across multiple experiments. This has a key implication when attempting to quantify the intensity of cell death caused by different treatments: comparing individual results across different leaves and plants is likely not a viable strategy (at least with this modality of generating HR), and instead, the averaging of multiple repeats is necessary to confidently establish differences. It would appear, however, that dose dependence holds when all samples are taken from the same leaf, and single experiments run in this format could provide at least a preliminary indication of cell death intensity.

**Figure 3.**
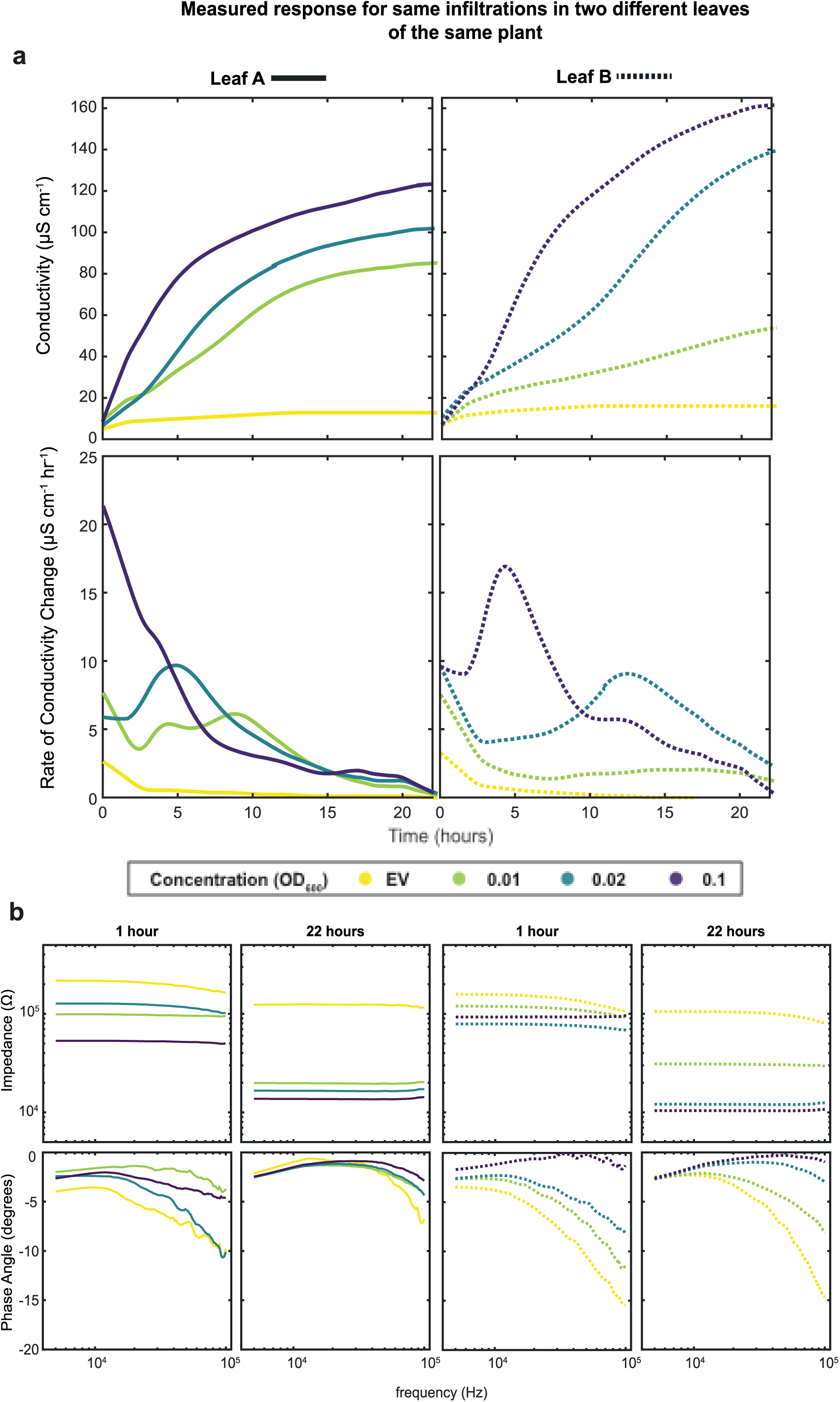
**a**) Conductivity (top) and rate of change of conductivity (bottom) plots for single electrolyte leakage experiment using same infiltrations on two leaves of the same plant (Left: Leaf A (solid), Right: Leaf B (dotted). Data captured at 10 kHz excitation frequency. LOWESS smoothing filter applied (span = 150). **b)** Measured impedance and phase angle of tested solutions plotted against frequency at time points of 1 and 22 hours, excitation frequency range 5 **–** 100 kHz. From left to right: (Leaf A: 1 hour and 22 hours, Leaf B: 1 hour and 22 hours)

Given the high temporal resolution of data generated by PASTEL, we were able to calculate the instantaneous derivative and rates of change of conductivity without the need for an interpolating numerical model. This allows the kinetics of cell death to be resolved with far more granularity than traditional hourly sampling, capturing differences in rates of change of conductivity that may otherwise be missed. For example, we observed secondary peaks in the rate of change of conductivity after the initial maxima from when discs were initially placed in the wells (**Figure 3b**). Maximal rates of change of conductivity appear to correlate with the concentration of infiltration (and so to the intensity of cell death), indicating this could be a useful additional metric for the quantification of HR (*i.e.* the rate of cell death).

To investigate the resistive and capacitive properties of our two-electrode electrochemical cell for studying HR (**Figure 3b**) we swept a range of frequencies (5 – 100 kHz) with 2 min intervals. Comparing data captured at 1 hour and 22 hours, we see the relationship between concentration of infiltration and solution impedance holds across the whole frequency span. Impedance measured relatively constant throughout the frequency range at both time points, with differences in magnitude between infiltrations decreasing slightly at higher excitation frequencies. Changes in solution impedance are not linearly correlated with the concentration of infiltration, and at the 22 hour end-point all samples tested lie within the 1.0×10^4^ – 2.0×10^4^ Ω band, with the exception of the negative control. On the other hand, we observe greater variation in phase angle with frequency, particularly at the high-frequency tail of the spectrum, where phase angle differs greatly depending on the concentration of infiltration.

Infiltrations resulting in higher impedance (lower conductance), in general, have more negative phase angles, indicating greater capacitance. Again, we observed variation between leaf A and leaf B for the same infiltrations; differences in impedance throughout the frequency range mirror the differences in conductivity seen in **Figure 3a**, but with the relationship of course inverted. Differences in phase angle are also seen, for example, phase angle is significantly more negative for the EV infiltration in leaf B than A at 22 hours.

In this perspective, the sole tracking of solution conductivity at a fixed frequency can obfuscate the determination of HR level. At low conductivities, low frequencies are favorable since the polarization resistance in 2-electrode configuration can be neglected. In contrast, the highly conductive solutions suffer from polarization resistance and necessitate a higher frequency for accurate determination of solution conductivity.^29^ The tracking of both impedance and phase angle on a frequency range can provide valuable insights as we can track a larger conductivity range accurately and provide a clearer picture of the level of HR in plant tissue.

### PASTEL enables detection and quantification of microscopic cell death

The ability to detect microscopic cell death is of great importance in plant immunity research; it facilitates the study of weak responses and early events occurring prior to HR symptoms becoming macroscopically visible (or in cases where they may not become visible at all). Detection of such low intensities of cell death requires highly sensitive techniques. In order to evaluate the performance of our system relative to an optical method of detection (which are known to be extremely sensitive), particularly with regard to very low intensities of cell death, we conducted electrolyte leakage and propidium iodide staining experiments simultaneously (**Figure 4**). Propidium iodide is a dye that is able to penetrate only non-viable cells and fluoresces upon intercalation with DNA.^30^ Cells with compromised membranes can be identified by imaging dyed tissue using fluorescence microscopy.

**Figure 4.**
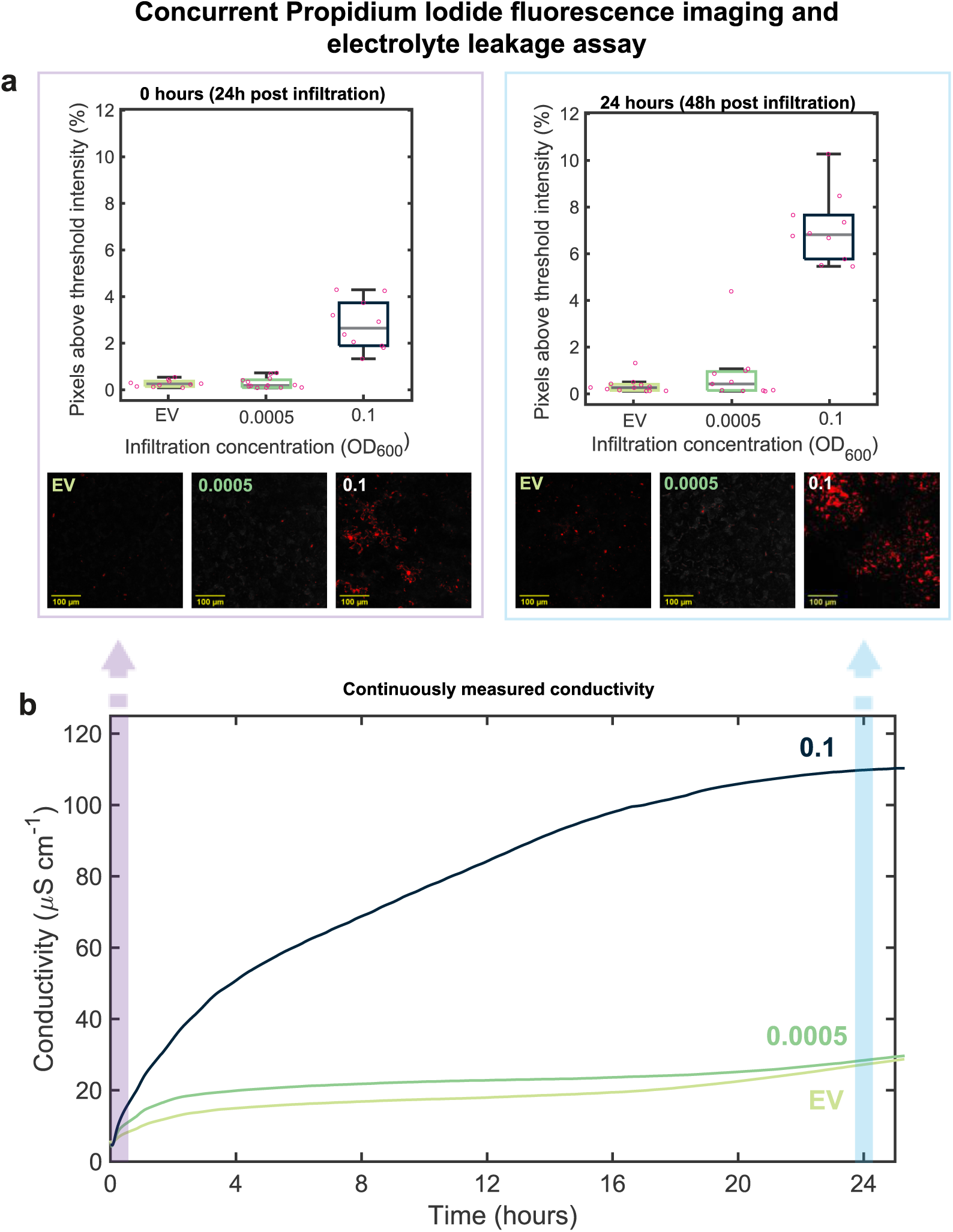
**a**) Box and whisker plot of fluorescence quantification performed on confocal microscopy imaging of propidium iodide stained leaf discs. Quantification was performed by projecting maximum intensity across the Z-stack and calculating the percentage of pixels above a threshold intensity value. Imaging was performed at the 0-hour timepoint (left) and 24-hour timepoint (right). Representative images following binary thresholding for each infiltration concentration below plots. **b)** Solution conductivity measured by our system during concurrent experiment performing an electrolyte leakage assay using discs from the same leaf. Data was captured at 10 kHz sampling frequency.

We excised leaf discs from the same leaf containing infiltrations to create negative and positive controls. We used OD_600_ = 0.0005 as the low concentration for microscopic scale cell death. For fluorescence imaging, discs were taken following 24 hours and 48 hours of incubation (corresponding to electrolyte leakage timepoints of 0 hours and 24 hours), stained and imaged. The fluorescence of each capture in the x-y plane was quantified by calculating the percentage of pixels above a fixed threshold intensity in an attempt to account for background fluorescence.

At 0 hours (24 hours post infiltration), we found the median percentage of pixels above the threshold intensity was significantly greater for the positive control (2.65) compared with the low concentration (0.19) and negative control (0.25) (**Figure 4a**). A difference in cell death intensity was not discernible with the low concentration and negative control at this time point, with the negative control in fact having a marginally greater median. In contrast, in our concurrent electrolyte leakage experiment we measured a higher conductivity for the low concentration relative to the negative control within an hour and throughout the experiment. In this instance, however, the difference in conductivity between the low concentration and the negative control decreased towards the endpoint (**Figure 4b**).

At 24 hours (48 hours post infiltration), we would expect greater cell death compared to the 0 hours measurement given the longer incubation period, and this is reflected in our results – we observed higher intensity fluorescence across a greater area of the leaf disc for the positive control, and a greater difference in median percentage of pixels above threshold intensity (6.82) versus the low concentration (0.42) and negative control (0.27). The median, mean, and maximum of the intensity measurement for the low concentration was now slightly greater than the negative control – this indicates resolving low intensities of cell death may be viable with this method of fluorescence-based quantification, although seemingly with a higher limit of detection than with our PASTEL platform.

### Machine learning models for binary and three-class classification

Utilizing our large dataset from replicate electrolyte leakage experiments conducted with *N. benthamiana*, we developed supervised machine learning models for detecting the onset of HR using the data acquired using PASTEL. We aimed to evaluate if application of machine learning models to our data could (i) aid in reliably detecting presence of HR for both low and high intensities of cell death (ii) classify different strengths of HR and (iii) reduce the required experimental run time to reliably determine presence of HR. We developed binary (No HR or HR) and three-class classification (No HR, Low HR or High HR) models.

We pre-processed our raw dataset by applying a calibration profile for gain-factor and phase-angle correction to produce calibrated conductance and phase data. Subsequently, we applied a time-domain Savitzky-Golay filter to remove noise from conductance and phase data captured at each frequency sampled. Each “run” (each time-series dataset from a single well) was assigned a unique ID and corresponding label for the target output vector. For binary classification, labels were assigned as “False” for negative control EV agroinfiltrations and “True” for all non-zero levels of agroinfiltration of AVRblb2-carrying suspensions. For three-class classification: “0” for negative control, “1” for low HR (0.0005 < OD_600_ **≤** 0.005) and “2” for high HR (0.01 **≤** OD_600_ **≤** 0.1). The classes were based on the visibility of HR seven days after agroinfiltration; infiltrations below OD_600_ = 0.01 showed little to no visible difference in comparison to the negative control and so were categorized as “low HR” (**Figure S3)**

To evaluate the performance of these approaches, we trained binary and three-class classification models at time intervals of 0–1 hours and 0–22 hours using corresponding subsets of the dataset. For example, for the 0–1 hour models, only data captured between those time points were used for training and testing. Time series-specific feature extraction was performed using the tsfresh library^31^ on conductance (magnitude, real and imaginary components), the time derivative of conductance and phase data captured at each frequency between 10 kHz – 100 kHz (10 kHz intervals). We trained each model using a random forest classifier and evaluated performance by ten-fold cross-validation, each fold comprised of training and test sizes of 80% and 20% of the input dataset, respectively. The pre-processing to model evaluation pipeline is summarized in **Figure S4**

We found the binary classification models performed well, with equally high cumulative true positive (TP) counts (“HR” correctly classified) with the 0–1 hour and 0–22 hours models (**Figure 5a**, **left**). With the 0–22 hours model, the true negative (TN) count (“No HR” correctly classified) was higher, leading to a higher mean accuracy (87.8%) than with the 0–1 hour model (84.1%) (**Figure 5b, left**), as defined in **Equation 1**. It should be noted that the TN counts were much lower than the TP counts owing to the inherently imbalanced nature of the experimental dataset (number of HR positive samples greatly exceeding that of negative controls). While the false positive (FP) count (“No HR” incorrectly classified as “HR”) was reduced with the 0–22 hour model, the false negative (FN) count (“HR” incorrectly classified as “No HR”) did not change. In other words, the precision (**Equation 2**) was improved, but the recall (**Equation 3**) was not. Owing to the improved precision, we obtained a higher F_1_ Score (**Equation 4**) with our 0–22 hours model (0.81) than with the 0–1 model (0.75). Additionally, the area under the curve (AUC) scores obtained from receiver operating characteristic (ROC) curves (**Figure 5c, left**) show the model using the 0–22 hours dataset has better classification performance (AUC = 0.81) than that with the 0-1 hour dataset (AUC = 0.73).

**Figure 5.**
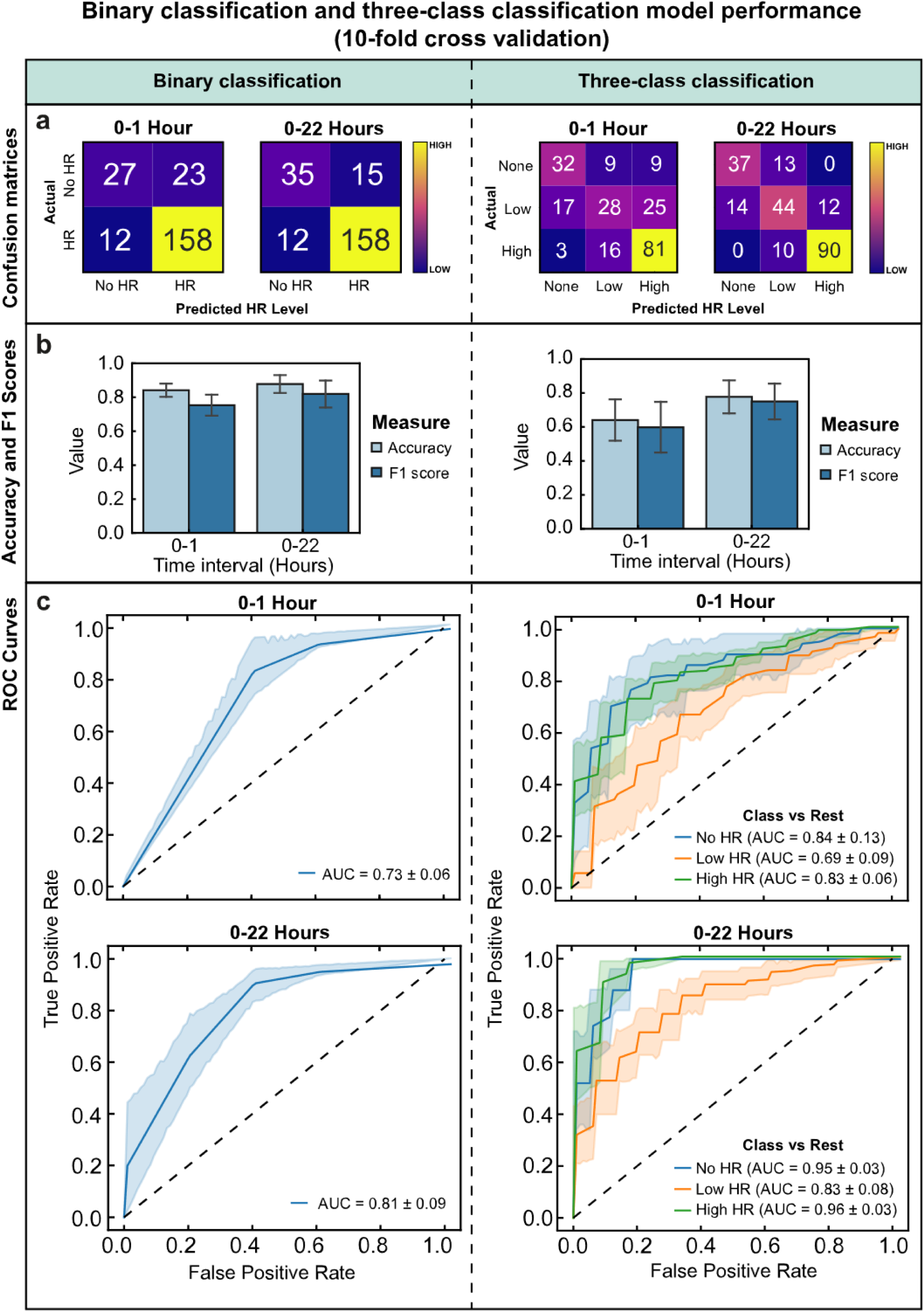
Left, all: performance metrics for binary (No HR/HR) classification models. **Right, all:** performance metrics for three-class (No HR/Low HR/High HR) classification models. 0–1 hours corresponds to models trained with the first hour of the experimental dataset and 0–22 hours corresponds to models trained with all 22 hours. **a)** Cumulative confusion matrices for 0–1 hour and 0–22 models. Actual vs predicted classes of tested samples summed across ten folds of cross validation. **b)** Accuracy and F_1_ scores for 0–1 hour and 0–22 models. Data represented as µ ± σ, with n = 10 folds of cross validation. **c)** Receiver operating characteristic curves of 0–1 hour and 0–22 hour models showing true positive rate against false positive rate at different classification thresholds. For the three-class classification models (right), ROC calculated as class vs rest. Legend indicates class and area under curve score. ROC and AUC data represented as µ ± σ, with n = 10 folds of cross validation.

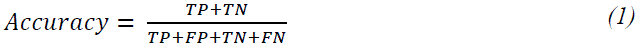

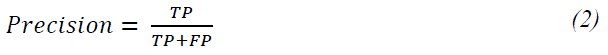

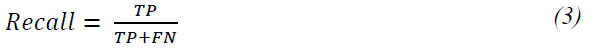

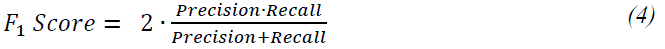

For three-class classification, there was a greater discrepancy in performance between the 0-1 hour and 0–22 hours models. With the 0–22 hours model, 30 additional samples were correctly classified, and unlike with the 0–1 hour model, there were zero misclassifications by more than one class across *i.e.* “No HR” misclassified as “High HR” or vice versa (**Figure 5a, right**). Accuracy and F_1_ improved significantly with the 0–22 hours model (Accuracy = 77.8%, F_1_ = 0.75) compared with the 0–1 hour model (Accuracy = 65.1%, F_1_ = 0.59) (**Figure 5b, right**). Expectedly, the one vs rest AUC scores (comparing each class against the rest) are also greater for the 0–22 hours model than the 0–1 hour model (**Figure 5c, right**). With the 0–22 hours model, AUC scores were particularly high for the “No HR” (AUC = 0.95) and “High HR” (AUC = 0.96) classes, indicating the model had a strong ability to discriminate these classes. “Low HR” had a lower score (AUC = 0.83), possibly due to the class neighboring the other two; the lowest concentrations within the class most at risk of being misclassified as “No HR” and highest most at risk of being misclassified as “High HR”.

In comparison to visual inspection of leaf tissue, our machine learning approach offers clear advantages in both detection time and sensitivity. At 1 day after infiltration (the time point at which leaf discs are excised) even the highest infiltration concentration (OD_600_ **=** 0.1) was not visually distinguishable from the negative control. Our binary-classification model was able to correctly predict HR with 84.1% accuracy one hour after this timepoint, using a test dataset inclusive of all concentrations tested. Moreover, at 2 days after infiltration visual symptoms of HR were only visible for concentrations of OD_600_ ≥ 0.02, while at 7 days visual symptoms remained absent for concentrations of OD_600_ **≤** 0.005.

## Discussion

Although PASTEL is a high-performance phenotyping system for quantifying HR cell death, it is constructed primarily from low-cost, off-the-shelf components, requiring only a computer for data recording and processing. The measurement wells are 3D-printed and reusable, while the disposable^32^ probes can quickly and easily be assembled by hand or using robots for large volume manufacturing. In any case, the PASTEL platform can be constructed in most academic settings with access to basic prototyping equipment. The cost of the entire system (∼$55) is substantially less than a research-grade benchtop conductivity meter that would typically be used in ELAs, with a consumable cost of ∼$0.53 per sample (**Table S1**). A simple, fast manufacturing process using low-cost, off-the-shelf components makes using a set of new sensors for each experiment viable, avoiding both the need for sterilization and the risk of electrode fouling.

Compared to traditional methods of manually sampling solution conductivity in a sequential manner, PASTEL significantly reduces both the time and labor necessary to conduct an ELA. In addition, it captures data at a much higher sampling rate (every 0–2 minutes versus the typical hourly rate). The high temporal resolution provides additional information about changes in conductivity that would otherwise be missed and enables rates of change in conductivity to be easily calculated. The excitation frequency is adjustable – measurements can be recorded at a single frequency or swept through a range at each time point, as opposed to typical fixed-frequency conductivity meters.

Automated sampling means no manual handling is required following the initial setup of an experiment, eliminating the potential risks of solution cross-contamination, disc perturbation, and data logging errors. Moreover, the large datasets generated from each experiment can be used to develop machine learning models for rapid binary and three-class classification of HR response with high accuracy, with the potential to reduce required experimental run time and automate the analysis of results. Use of machine learning with PASTEL has potential applications in high throughput phenotyping; both minimizing the required experimental run-time and negating the need for manual data analysis to determine assay results.

The PASTEL platform has the following disadvantages: (i) As it is based on the ELA protocol, it suffers from the same sample preparation drawbacks; disc excision is time-consuming and requires careful handling, a wash step is required, and the process is destructive to the plant.

Nevertheless, comparatively less handling is required than most other HR assays with the exception of whole-leaf fluorescence imaging. (ii) A negative control is needed as a reference point to determine whether a sample is undergoing HR, as basal electrolyte leakage can vary. Use of a machine learning model generated from previous experiments with the same plant species could remove this requirement. (iii) Conductivity measurements are highly temperature sensitive and so the system must be run in a temperature-controlled environment, especially if comparing separate experiments.

Additional calibration protocols and temperature sensors may also be used for temperature compensation.

PASTEL has high scalability and could be evolved into a high-throughput rapid phenotyping platform simply by integrating a larger multiplexer and additional well arrays. Through electrode functionalization, it could also be further developed toward a selective ion sensing approach aiming to temporally resolve individual ion species released into the analytical media in the HR. This development path could shed light on the complex mechanisms of ETI-based HR response and intercellular signaling, providing an invaluable and versatile tool for plant science. While in this work, we only evaluated the system with *N. benthamiana* undergoing HR, PASTEL could be adapted for use with other species, other forms of cell death or indeed any response which would result in a sufficiently high flux of electrolytes from tissue to solution.

## Methods

### Experiments

#### Plant Material

Transgenic *Nicotiana benthamiana* expressing Rpi-blb2 were soil-grown in a growth chamber with a day/night cycle of 24°C, 50% RH (relative humidity) and 22°C, 65% RH respectively. The day length was set at 16 hours. 4–5 weeks old plants were used for experiments. For each individual experiment, two leaves from a singular plant were used.

#### Transient gene expression

*Agrobacterium tumefaciens* carrying AVRblb2-3xHA constructs^33^ were used to mediate transient expression of AVRblb2 in Rpi-blb2-transgenic *N. benthamiana* leaves and thereby generate HR.^24,26^ *A. tumefaciens* carrying empty vector EV-3xHA constructs^34^ were used as control. *A. tumefaciens* cultures were grown for 2 days on LB agar plates with spectinomycin selection (Tryptone 10 g/L, Yeast Extract 5 g/L, Sodium Chloride 10 g/L, Agar 15 g/L, spectinomycin 100 µg/mL. Washing and harvesting: bacteria was twice suspended in 1.5 mL sterile dH_2_O and centrifuged at 4000 rpm for 5 minutes before being resuspended in 1 mL agroinfiltration buffer (10 mM MES hydrate, 10 mM MgCl_2_). The optical density of the stock suspension at 600 nm was measured with a spectrophotometer (Eppendorf) and then diluted to the desired OD_600_ value. In dose-dependent tests, AVRblb2 construct-carrying suspensions and empty vector construct-carrying suspensions were mixed to obtain an overall OD_600_ of 0.1. Inoculation of the plant material was performed by pressure-infiltration with a needleless syringe on the abaxial leaf surface on whole plants.

#### Assay procedure

*A. tumefaciens* suspensions were prepared at the desired OD_600_ values, and pressure infiltrated into leaves of Rpi-blb2-transgenic *N. benthamiana*. Infiltrated regions were outlined with a marker for identification when dry. Following a 24-hour incubation of the plant in a growth room, leaf discs were excised using a number 3 cork borer (7.8 mm diameter) in sets of 3, with each set then placed in a 1.5 mL centrifuge tube containing 1ml sterile dH_2_O for 15 minutes. The tubes were inverted several times before transferring each set of discs to a test well (1 set per well) containing 2 mL sterile dH_2_O. Lids were attached to the test wells, and measurement was initiated by serial command on the connected computer. Following the measurement period, data was imported and processed in MATLAB. Experiments were conducted in a temperature-controlled incubator maintained at 24°C.

#### Propidium Iodide Staining and Confocal Microscopy

20 µM Propidium iodide (PI) solution was prepared by dilution of 1.0 mg·mL^-1^ stock (Merck). Leaf discs were excised from the treated plant and placed in separate centrifuge tubes containing 20 µM PI to incubate for 15 minutes. Discs were removed, placed in a petri dish with dH_2_O and then washed for 5 minutes using an orbital shaker. Microscopy slides were then prepared with the discs for imaging with a confocal microscope using a water immersion lens. 10 captures were taken across the x-y plane per leaf disc, with 6-9 z-axis slices taken per capture. Data were processed and analyzed using ImageJ (Fiji) to quantify the fluorescence intensity. For each capture, a maximum intensity Z-axis projection was applied. Binary thresholding was applied to the resulting image (< 250 → 0, ≥ 250 →1) in order to account for background fluorescence. The mean intensity of the binary image was then quantified and normalized to give a percentage of pixels above the threshold intensity.

### System

#### Modular wells

The modular wells were 3D printed (N2 Plus, Raise3D) with transparent ABS filament (Verbatim). The lids and the well holder were 3D printed with orange PLA filament (Raise3D). All printed components were designed using SOLIDWORKS.

#### Conductivity Probes

Conductivity probes were constructed by embedding two-pin gold plated through the hole connector header (5-826634-0, TE Connectivity) into the lid of a 1.5 mL reaction tube (616-201, Greiner Bio-one). Pins were aligned with the lid brim and attached to jumper connectors (PRT-12794, SparkFun Electronics). The assembly was secured in position using hot melt adhesive. A 6.5 mm bore nitrile rubber O-Ring (RS PRO) was placed around the lid aperture to create a seal when inserted into the modular well.

#### Calibration

Conductivity calibration of the system was performed using dH_2_O and KCl solutions ranging from 2 µM – 50 mM. Calibration solutions were kept inside a temperature-controlled incubator set to 24°C prior to testing. For each solution tested, 2 mL was placed in an individual well and conductivity was measured for 30 minutes (frequency sweep 2 – 100 kHz, 2-minute sweep interval) and repeated (n = 3). Wells were sterilized between measurements, and new electrode assemblies were used. A research grade pH/Conductivity bench meter was used for reference conductivity measurements of the same solutions (n = 3) (HI-5521, HANNA Instruments).

Impedance magnitude and phase angle data were calculated from AD5933 Impedance converter’s raw output, as outlined in the datasheet,^35^ for use in the machine learning model. To correct system offset and gain errors in measured magnitude and phase angle, calibration data was obtained by performing a frequency sweep on an experimentally relevant range of resistive loads (1 – 220 kΩ). For gain factor correction, a mapping of measured magnitude to actual impedance was generated by finding coefficients for a 4th-order polynomial fitting at each excitation frequency (5 – 100 kHz). For phase angle correction, a mapping of measured magnitude to the system phase angle was generated by finding coefficients for a two-term exponential fitting at each excitation frequency. The mappings were then applied to the raw data, converting magnitude to impedance and subtracting the calculated system phase angle from the measured phase angle.

For experimental data captured at the 10 kHz excitation frequency, raw output magnitude values were converted to conductivity using a scaling factor calculated from the average calibration measurements of 500 µM KCl. Constant sensor parameters and a constant gain factor throughout the relevant range of impedance at this frequency allowed for simple linear scaling.^36–39^

#### Data Acquisition and Processing

Measurement parameter input, measurement initiation and data capture were performed using SerialPlot (Hasan Yavuz) interfacing with the ATmega328P CH340 Nano microcontroller. The microcontroller was programmed using the Arduino IDE. MATLAB (Mathworks) was used to process the comma-separated value (CSV) output files from each experiment, apply calibration profiles and perform data analysis. Overall circuit schematic shown in **Figure S6.**

### Machine Learning Model

The magnitude and phase-corrected phase and conductance data were pre-processed by time-domain Savitzky-Golay filtering (span = 50% window size) at each measured frequency. Each data series from each individual well across all experiments was assigned an ID and label for the target output vector (False if EV, True otherwise). Feature extraction was performed using tsfresh library (Input variables: conductance, phase, rate of change of conductance at each measured frequency).^31,40^ Models were trained by random forest classification (sklearn) using the balanced class-weighting parameter and cross-validated using stratified shuffle split (sklearn) with 10 splits and an 80:20 training: test ratio.^41^ Accuracy and F1 scores were recorded for each fold.

