## Supplementary Information for "Parallel, Continuous Monitoring and Quantification of Programmed Cell Death in Plant Tissue"

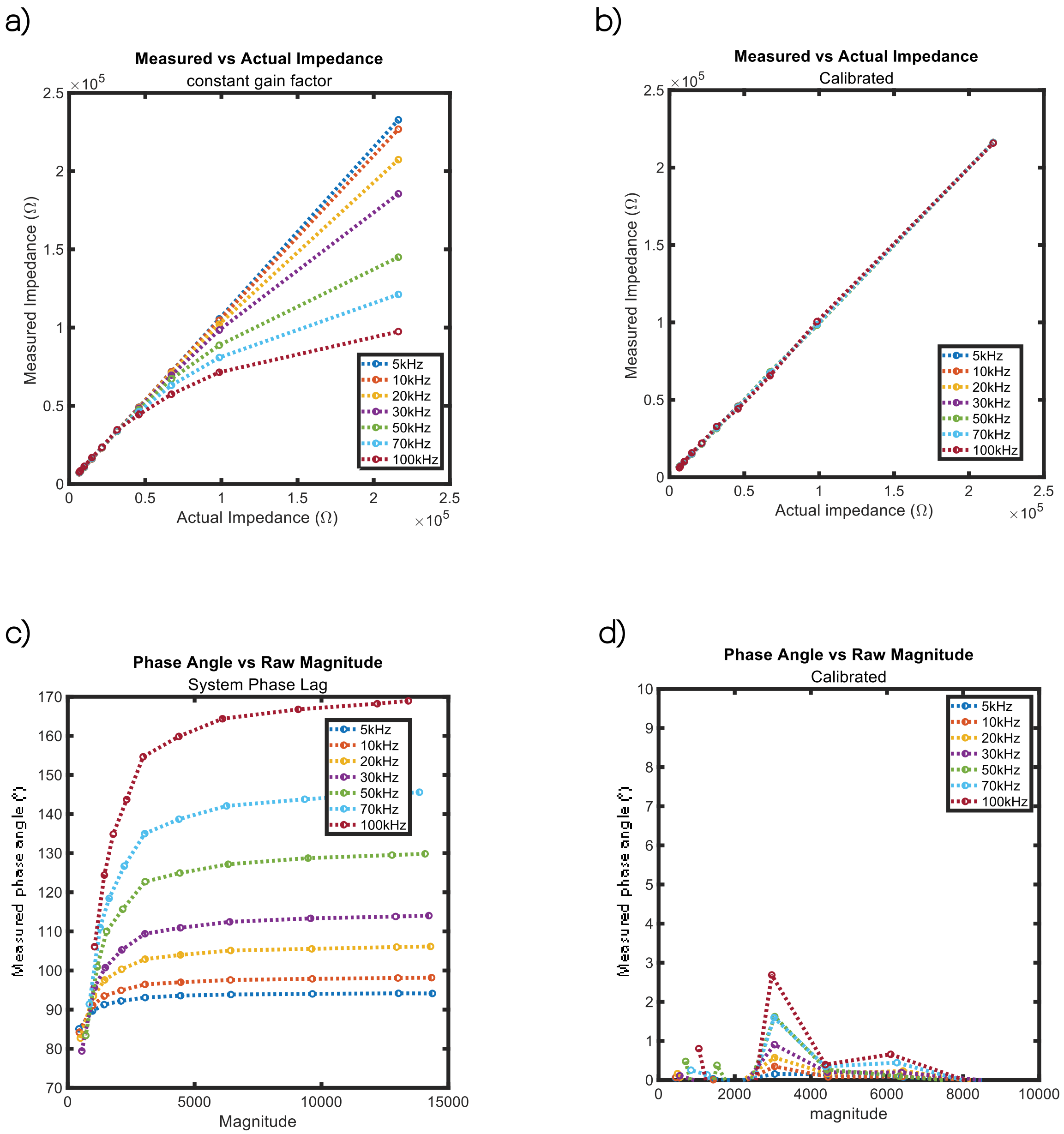


**Figure S1. Gain factor and Phase Angle System Calibration: a)** Correlation plot of measured and actual impedance, measuring resistances in the range 6.6-220 kΩ using the PASTEL system. Impedance calculated using constant gain factor calculated as described in **Equation S2 and Equation S3. b)** Correlation plot of measured and actual impedance, measuring resistances in the range 6.6-220 kΩ using the PASTEL system. Impedance calculated using 4$^{th}$ order polynomial mapping function (**Equation S4**), derived from application of a fitting function performed on data obtained in a). **c)** Correlation plot of system phase angle against raw magnitude measured, measuring resistances in the range 6.6-220 kΩ using the PASTEL system. **d)** Correlation plot of measured phase angle against raw magnitude measured, measuring resistances in the range 6.6-220 kΩ using the PASTEL system. Measured Phase angle calculated as described in **Equation S5** using a power mapping function derived from application of a fitting function performed on data obtained in c).

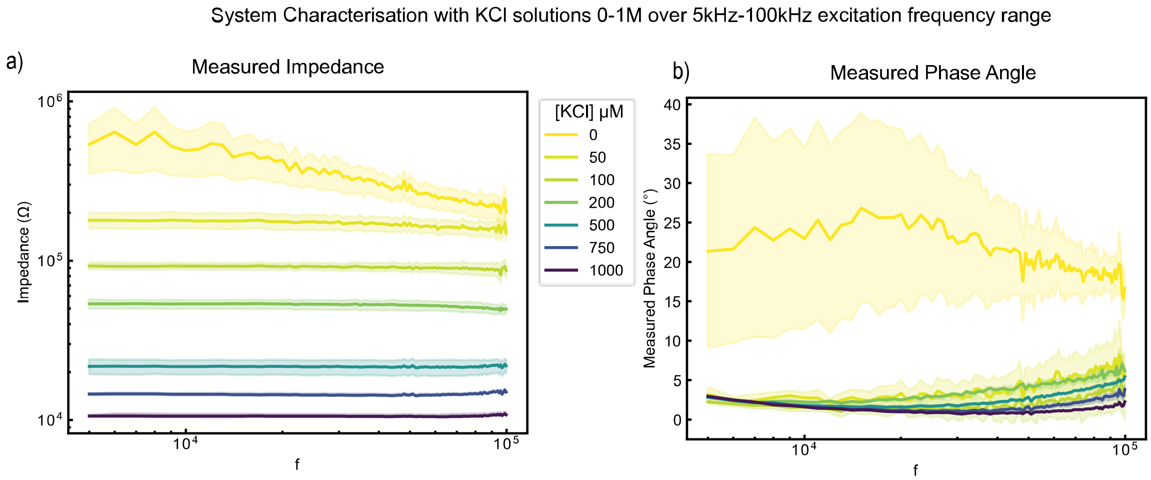


**Figure S2. System Characterization with Potassium Chloride (KCl)**: **a)** Impedance measured with PASTEL using range of KCl concentrations (0 — 1.0 M) against excitation frequency Data captured at 20 minutes. Data represented as µ ± $\sigma$, with n = 3 independent samples **b)** Phase angle measured with PASTEL using range of KCl concentrations (0 — 1.0 M) against excitation frequency Data captured at 20 minutes. Data represented as µ ± $\sigma$, with n = 3 independent samples.

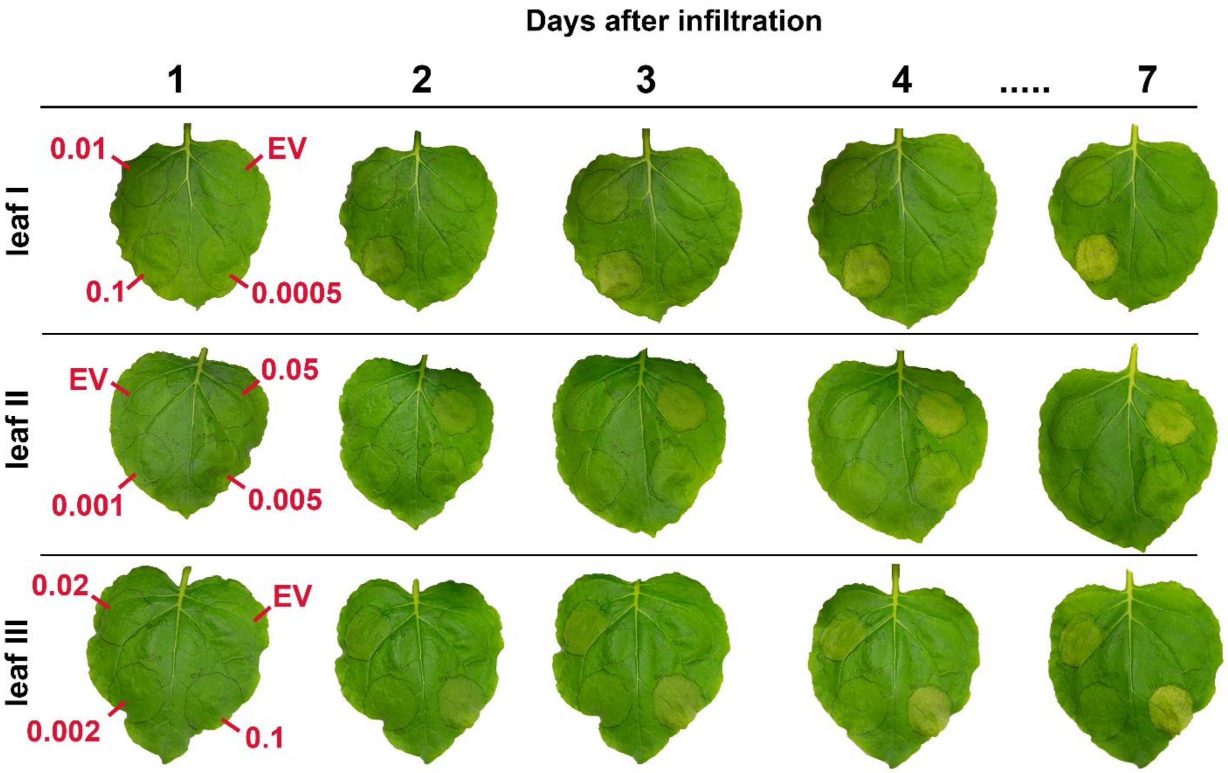


**Figure S3. Visual HR Symptoms at 1-4 and 7 days after infiltration.** Images of leaves agroinfiltrated with an empty vector-carrying bacterial suspension (negative control) and AVRblb2-carrying bacterial suspensions of concentrations in the range $OD_{600}$ = 0.0005-0.1.

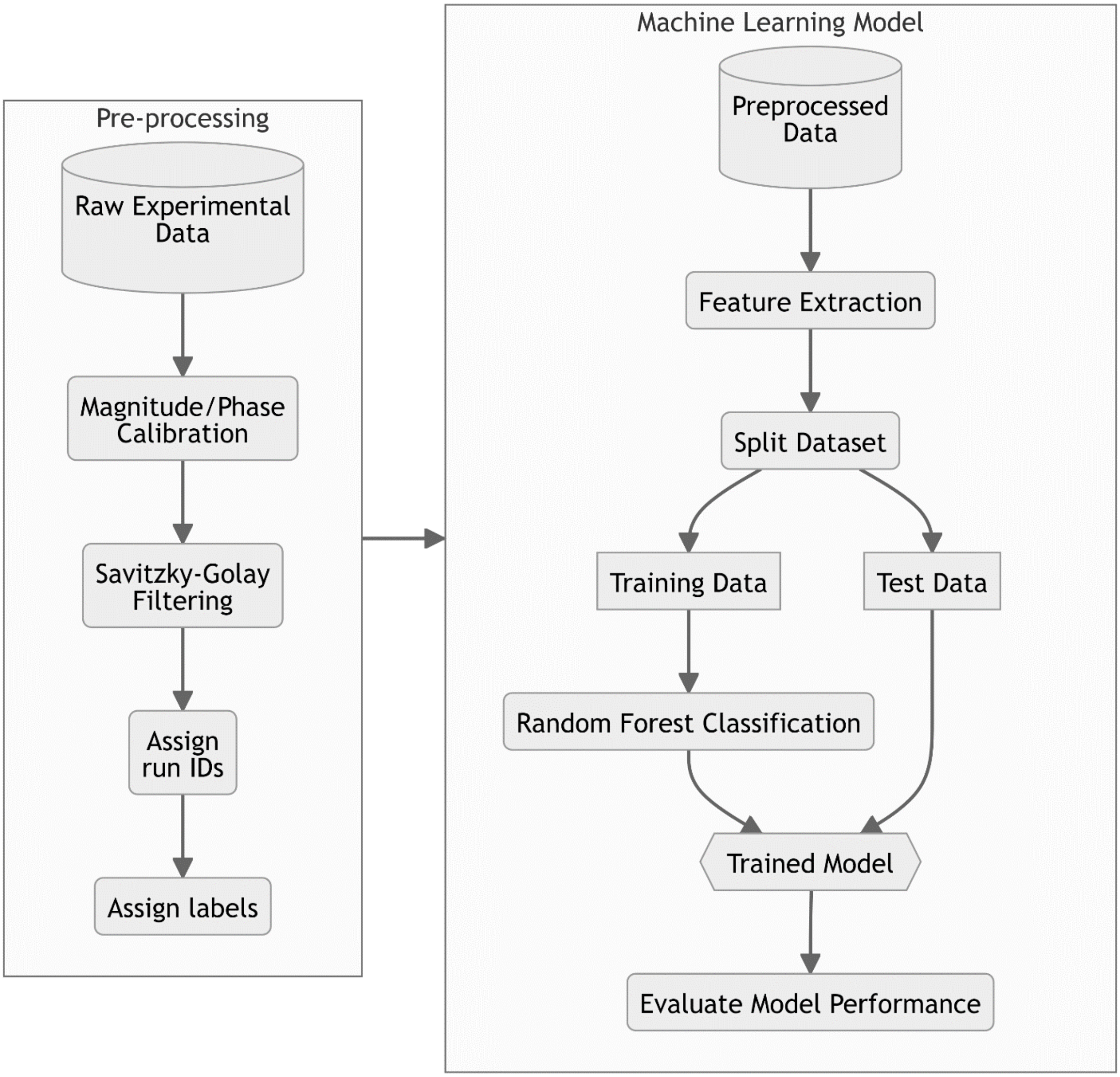


**Figure S4. Flowchart outlining data pre-preprocessing steps and development of machine learning models for classification of HR**

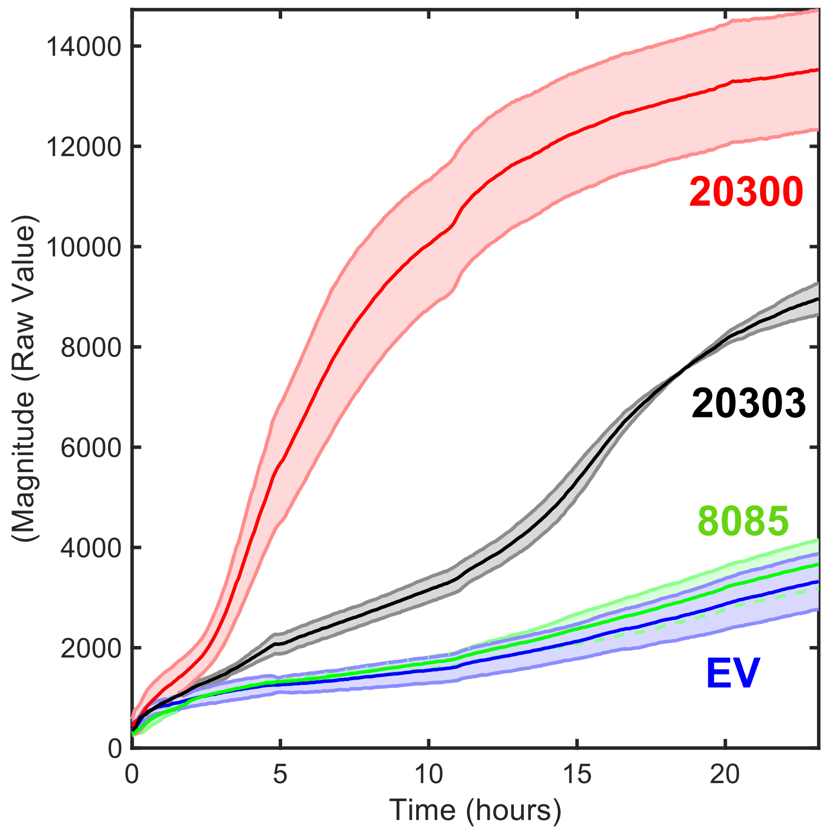


**Figure S5. Electrolyte leakage experiment with AVRblb2 paralogs conducted with an earlier version of PASTEL.** A leaf of Rpi-blb2 transgenic *N.benthamiana* was infiltrated with suspensions carrying constructs of EV and AVRblb2 paralogs PITG_20300, PITG_20303 and PITG_8085 at $OD_{600}$ = 0.1. PITG_20300 is the reference paralog used in all other experiments. One leaf disc was placed in each well (an Eppendorf tube) in 1ml $dH_2O$. Data n=2, shaded region represents max and minimum values.

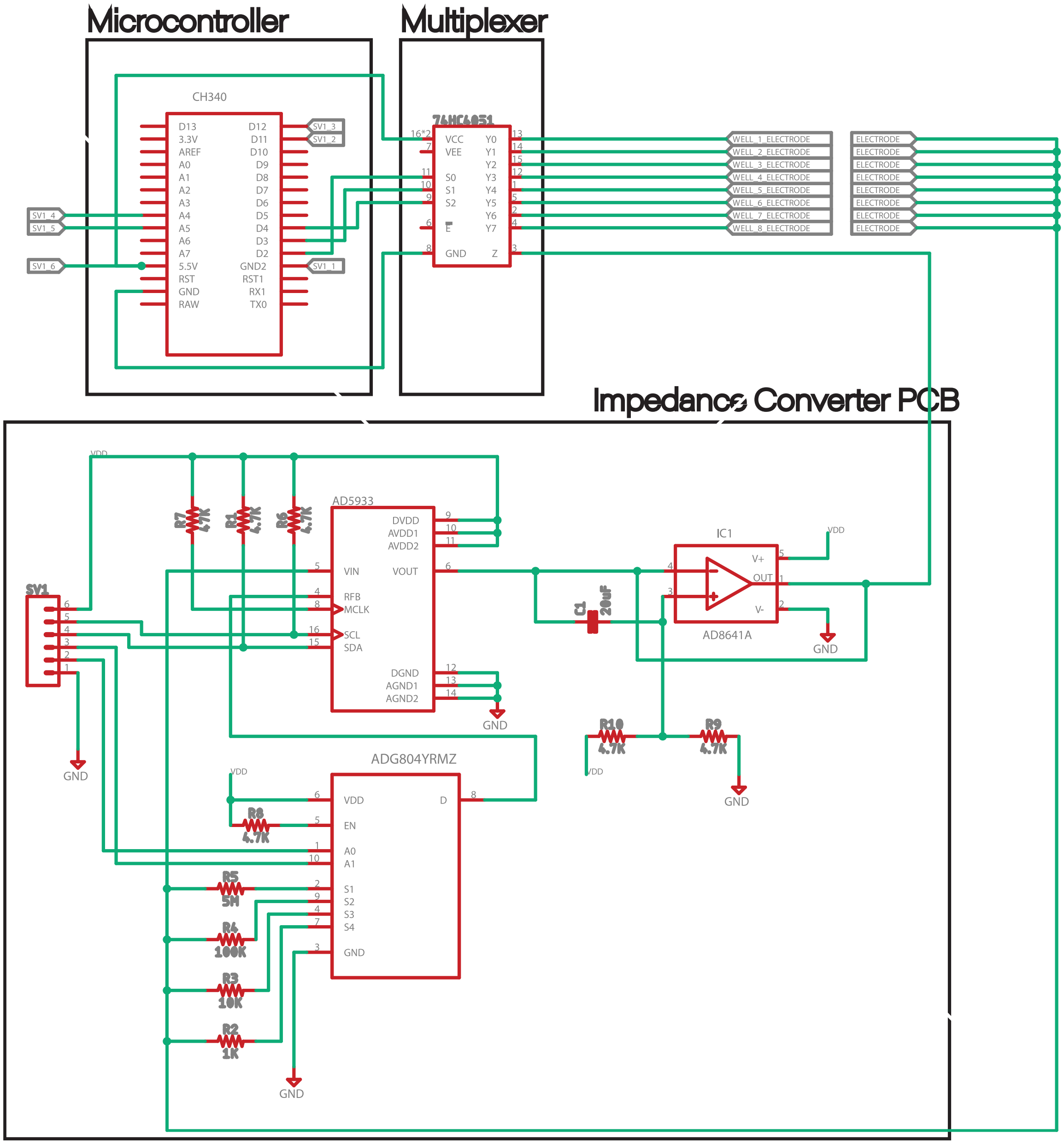


**Figure S6.** System electronics schematic of microcontroller, multiplexer and custom printed circuit board.

**Equation S1: Conversion of Raw Magnitude (at 10kHz) to Conductivity**

$$\sigma = \gamma \, M_{PASTEL}$$

$$\gamma = \bar{\sigma}_{HANNA(500\,\mu M\,KCl)} / \bar{M}_{PASTEL(500\mu M\,KCl)}$$

$$\gamma = 85.7 \,/\, 4869.32$$
$$\gamma = 0.0176\ \mu S \cdot cm^{-1}$$

where $\gamma$ = scale factor, $\sigma$ = conductivity and $M_{PASTEL}$ = raw magnitude measured by PASTEL.

Constant scale factor can be used at for measurements performed with 10 kHz excitation frequency as system measurements are linear throughout the relevant impedance range at this frequency (**Figure S2a**).

**Equation S2: Recommended Calibration Resistance**[1]

$$R_{CAL} = \frac{Z_{min}+Z_{max}}{3}$$

$$75.5k = (220k + 6.6k)/3$$

$$\rightarrow\ 67\ k\Omega$$

**Equation S3: Constant Gain Factor Calculation**[2]

$$Gain\ Factor = Magnitude_{50kHz}/R_{CAL}$$

$$\text{Gain Factor} = 1164665.86/67000$$

$$Gain\ Factor = 9.70579e - 9\ \Omega^{-1}$$

$$Magnitude = \sqrt{(R^2 + I^2)}$$

where R = Real and I = Imaginary data word outputs from the AD5933 Impedance Converter.

**Equation S4: Impedance calibration**[2]

$$Z(x,f) = a_f x^3 + b_f x^2 + c_f x + d_f$$

where $Z$ is the calibrated impedance, $f$ is frequency, $x = \frac{Gain\ Factor}{Raw\ Magnitude}$, and $a_f$, $b_f$, $c_f$, $d_f$ are frequency dependent constants.

**Equation S5: Phase Angle calibration**[2]

$$Z\emptyset\ =\ \Phi\ -\ \nabla$$

where $Z\emptyset$ is the phase of the unknown impedance, $\Phi$ is the raw phase angle and $\nabla$ is the system phase angle

$$\nabla(M, f) = a_f e^{b_f M} + c_f e^{d_f M}$$

where $\nabla$ is the system phase angle, $M$ is the raw magnitude, and $a_f, b_f, c_f, d_f$ are frequency dependent constants.

**Table S1. Table showing the breakdown of system cost and consumables per experiment sample.** All prices are in USD.

| | | |
|---|---|---|
| **System** | | |
| **Component** | **Supplier** | **Price (USD)** |
| ATmega328P CH340 Nano | Kunkune | 6.27 |
| AD5933YRSZ | Digikey | 22.69 |
| AD8641A | Digikey | 4.98 |
| ADG804YRMZ | Digikey | 4.66 |
| Sparkfun 74HC4051 | Pimoroni | 3.46 |
| PCB Manufacturing + SMD Components | Elecrow | 6.4 |
| ABS Filament (8 Wells) | Verbatim | ~3.84 |
| PLA Filament (Well Holder) | RS | ~2.56 |
| **Total** | | **~54.86** |
| **Consumables (per well)** | | |
| Jumper PRT-12794, SparkFun Electronics | Digikey | 0.21 |
| 616-201, Greiner Bio-one | Greiner Bio-one | ~0.03 |
| TE Connectivity 5-826634-0 | Mouser | 0.23 |
| RS PRO Nitrile Rubber O-Ring, 6.5mm Bore, 10.5mm Outer Diameter | RS | 0.06 |
| **Total** | | **~0.53** |

### Supplementary References

[1] Analog Devices Inc., AN-1252 Technical Note, http://www.analog.com/media/en/technical-documentation/application-notes/AN-1252.pdf

[2] Analog Devices Inc., AD5933 Datasheet, https://www.analog.com/media/en/technical-documentation/data-sheets/AD5933.pdf
